# Context dependent contributions of the direct and indirect pathways in the associative and sensorimotor striatum

**DOI:** 10.1101/2024.01.18.576337

**Authors:** Nisa Cuevas, Argelia Llanos-Moreno, Kathia I. Ramírez-Armenta, Hector Alatriste-León, Josué O. Ramírez Jarquin, Fatuel Tecuapetla

**Author notes:** For correspondence (FT). These authors contributed equally to this work (Nisa Cuevas, Argelia Llanos-Moreno). Ernst Strüngmann Institut, Faculty of Biological Sciences, Goethe Universität, and International Max Planck Research School for Neural Circuits, Frankfurt am Main, Germany (Nisa Cuevas). Instituto de Neurobiologia, UNAM, México. Psychology Department, University of California, Los Angeles, U.S.A.

## Abstract

To determine whether the contributions of striatal projection neurons from the direct (dSPNs) and indirect (iSPNs) pathways of the basal ganglia to action selection and locomotion can be generalized across the associative (DMS) and sensorimotor (DLS) striatum we compared the optogenetic activation or inhibition of these pathways on different tests. We show that self-modulation of dSPNs or iSPNs in either compartment has opposite contributions to real-time place preference, and to selecting an action in the DMS but not in the DLS. During reward seeking displacements, activation of either pathway in both compartments, or inhibition of dSPNs in the DMS slows movement. During spontaneous displacements, dSPNs activation showed opposing effects depending on the compartment modulated. Remarkably, inhibition of either pathway in the DLS decreases while only iSPNs inhibition in the DMS facilitates these displacements. These findings support a model of opposite, complementary and undescribed contributions of the striatal pathways depending on the compartment and context.

## Introduction

Basal ganglia (BG) circuits are involved in action selection (Grillner et al., 2005; Gurney et al., 2001; Hikosaka et al., 2006; Humphries et al., 2006; Mink, 1996; Redgrave et al., 1999) and control of movement (Rubchinsky et al., 2003; Kravitz et al., 2010; Tecuapetla et al., 2016). The striatum the largest nuclei of the BG receives glutamatergic input from the cortex and thalamus, as well as dopaminergic and serotoninergic input from the brain stem (Redgrave et al., 1999). Striatal spiny projection neurons (SPNs) are GABAergic and are segregated into two pathways. Indirect pathway neurons (iSPNs) project to the external globus pallidus and express D2-type dopamine receptors and adenosine A2A receptors, whereas direct pathway neurons (dSPNs) project to the output nuclei of the BG, and express D1-type dopamine receptors (Gerfen et al., 1990; Schiffmann and Vanderhaeghen, 1993; Kita and Kitai, 1988). BG loops can be differentiated as limbic, associative, and sensorimotor, and are anatomically identified in rodents as the ventral striatum (VS), dorsomedial striatum (DMS), and dorsolateral striatum (DLS) respectively (Alexander et al., 1986; Draganski et al., 2008). Lesions in the DMS interfere with the capacity to perform goal-directed actions (Yin et al., 2005a,b) while lesions in the DLS disrupt habit formation (Yin et al., 2004, 2006). The rate model of the basal ganglia pathways proposes that activation of dSPNs promotes movement and activation of iSPNs inhibits movement (Albin et al., 1989; DeLong, 1990; Penney and Young, 1983). Evidence supporting this model was provided by optogenetic activation of these pathways in the DMS (Kravitz et al., 2010; Isett et al., 2023). Furthermore, operant self-stimulation of dSPNs in the DMS induces reinforcement, but in iSPNs triggers avoidance (Kravitz et al., 2012). In contrast to this model, other studies performed in the dorsal striatum or DLS have showed co-activation of both dSPNs and iSPNs during the initiation of actions, suggesting that the pathways act cooperatively. (Cui et al., 2013; Jin et al., 2014; Tecuapetla et al., 2016). Frequently basal ganglia circuits, specifically the striatum, are referred to as a unit despite evidence showing differential contributions of the striatal compartments to reinforcement-guided action (Yin et al., 2005b,a, 2004, 2006). To date it is still unclear whether the contributions of projection neurons from the direct and indirect pathways of the basal ganglia, to action selection and locomotion, can be generalized across the associative (DMS) and sensorimotor (DLS) compartments of the striatum. Previous work supporting either the rate model or a complementary contribution of the pathways has focused on either activating the striatal pathways, or measuring their activity in specific compartments, without accessing whether these activations or measurements can be generalized across the DMS and DLS (Kravitz et al., 2010, 2012; Cui et al., 2013; Jin et al., 2014). Importantly, experiments investigating inhibition of the activity of cells in these pathways are also lacking. This has left an open question: Can the contribution of striatal pathways be generalized between DMS and DLS? One possibility is that the contribution of striatal cells depends on the anatomical compartment and/or on the demands of the context, since each compartment receives different inputs. To evaluate this, we used optogenetics to activate or inhibit dSPNs or iSPNs in the DLS versus the DMS during three behavioral different tests.

## Results

To test whether the contribution of the striatal pathways can be generalized between the DMS and DLS in different contexts we performed optogenetic activation or inhibition of dSPNs or iSPNs in the DMS versus the DLS in three conditions 1) in a real-time place preference test (RT-PP), 2) in an action selection test, and 3) during reward seeking or spontaneous displacement. We and other groups have previously documented the feasibility of expressing ChR2 or Arch3.0 in SPNs to activate or inhibit these neurons selectively in the DLS (Tecuapetla et al. 2014, 2016) or the DMS (Kravitz et al., 2010; Tai et al., 2012; Ramírez-Armenta et al., 2022).

### Optogenetic self-stimulation of dSPNs or iSPNs in the DMS or DLS supports opposing contributions to real-time place preference

It has been previously reported that self-stimulation of striatal dSPNs or iSPNs in the DMS evokes approaching and avoidance respectively (Kravitz et al. 2012), however a direct comparison between the DLS and DMS has not been performed. We evaluated the effects of self-activating dSPNs or iSPNs in the DMS versus the DLS in a real-time place preference test (RT-PP) [animals received 20 Hz, 10 ms pulses, as long as they remained in a virtually selected quadrant of an open field; ***Figure 1A***]. Animals that received self-stimulation in dSPNs, either in the DMS or DLS, spent more time in the quadrant paired with self-stimulation [mean±SEM, preference index DMS-D1-ChR2=0.66 ±0.04, (n=9), DLS-D1-ChR2=0.57±0.06, (n=7), Controls=0.05±0.06, (n=12; 6 DMS and 6 DLS), p<0.001, Kruskal Wallis test, ***Figure 1B***, upper panel and ***1C***]. On the contrary, animals that received self-stimulation in iSPNs avoided the stimulation quadrant [mean±SEM, preference index DMS-A2A-ChR2=-0.73±0.07, (n=5), DLS-A2A-ChR2=-0.79±0.04, (n=7), Controls=-0.22±0.14 (n=8), p<0.002, Kruskal Wallis test, ***Figure 1B***, bottom panel and ***1D***].

**Figure 1.**
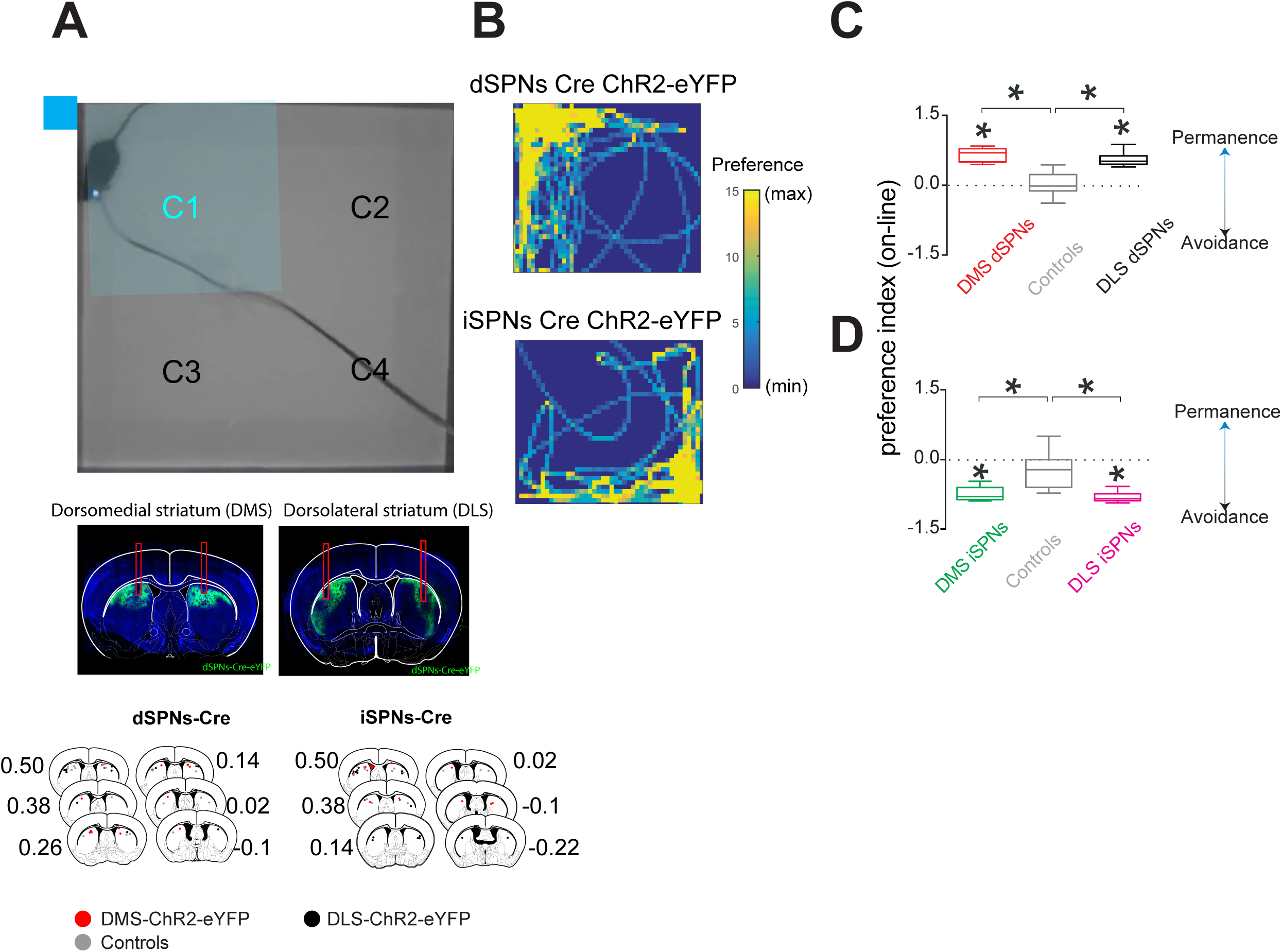
Optogenetic activation of dSPNs or iSPNs evokes preference or avoidance respectively in the real-time place preference test. **A**. An open field was virtually divided into four quadrants, the quadrant where an animal received self-stimulation is highlighted in blue. Middle panels show representative photomicrographs of coronal brain slices from animals with bilateral DMS or DLS implants. Bottom panels, schematic representation of the position of the fiber’s tips. **B**. Representative examples of the trajectories of two different animals, color coding the time spent per pixel during the test. Activation of dSPNs or iSPN are shown in the upper and lower panels respectively. **C**. Preference index of the time that animals spent in the quadrant with self-stimulation compared to an average of the other three quadrants for dSPN activation. **D**. Same as C, but for self-activation of iSPNs. In C and D, * above the horizontal lines is the Dunńs multiple comparison test p<0.001, post Kruskal Wallis. * above a group is p<0.001 Wilcoxon test.

To obtain a clearer picture of the contribution of the two pathways we performed the same experiment with self-inhibition i.e., animals received continuous light inhibition as long as they remained in a virtually selected quadrant of the open field, ***Figure 2***. Animals in which dSPNs were self-inhibited, either in the DMS or DLS, did not display a significant preference compared to controls when averaging across the 20 minutes of the test (***Figure 2C***). However, segmenting the total test time into 5 minute bins showed that animals developed avoidance to the inhibition quadrant, with the 10-15 minute bin being significantly different to that of the control group [mean±SEM, preference index DMS-D1-Arch3.0=-0.32±0.07, (n=7; 2 D1-Cre/Ai35), DLS-D1-Arch3.0=-0.16±07, (n=8; 2 D1-Cre/Ai35), Controls=0.05±0.03, (n=13), p<0.05, Mann Whitney U test, ***Figure 2D***]. The opposite was observed when inhibiting the iSPNs, either in the DMS or the DLS, either when averaging across the 20 minutes test time (Kruskal Wallis test, ***Figure 2E***), or during the last 10 minutes of the test [mean±SEM, preference index DMS-A2A-Arch3.0=0.15±0.09, (n=10; 3 A2A-Cre/Ai35), DLS-A2A-Arch3.0=0.30±0.13, (n=13; 4 A2A-Cre/Ai35), Controls=0.01±0.02, (n=13), p<0.05, Mann Whitney U test, ***Figure 2F***). Together these results show that self-modulation of the striatal pathways during real-time place preference yields opposite effects in both striatal compartments, with dSPNs supporting preference and iSPNs avoidance behaviors in this test.

**Figure 2.**
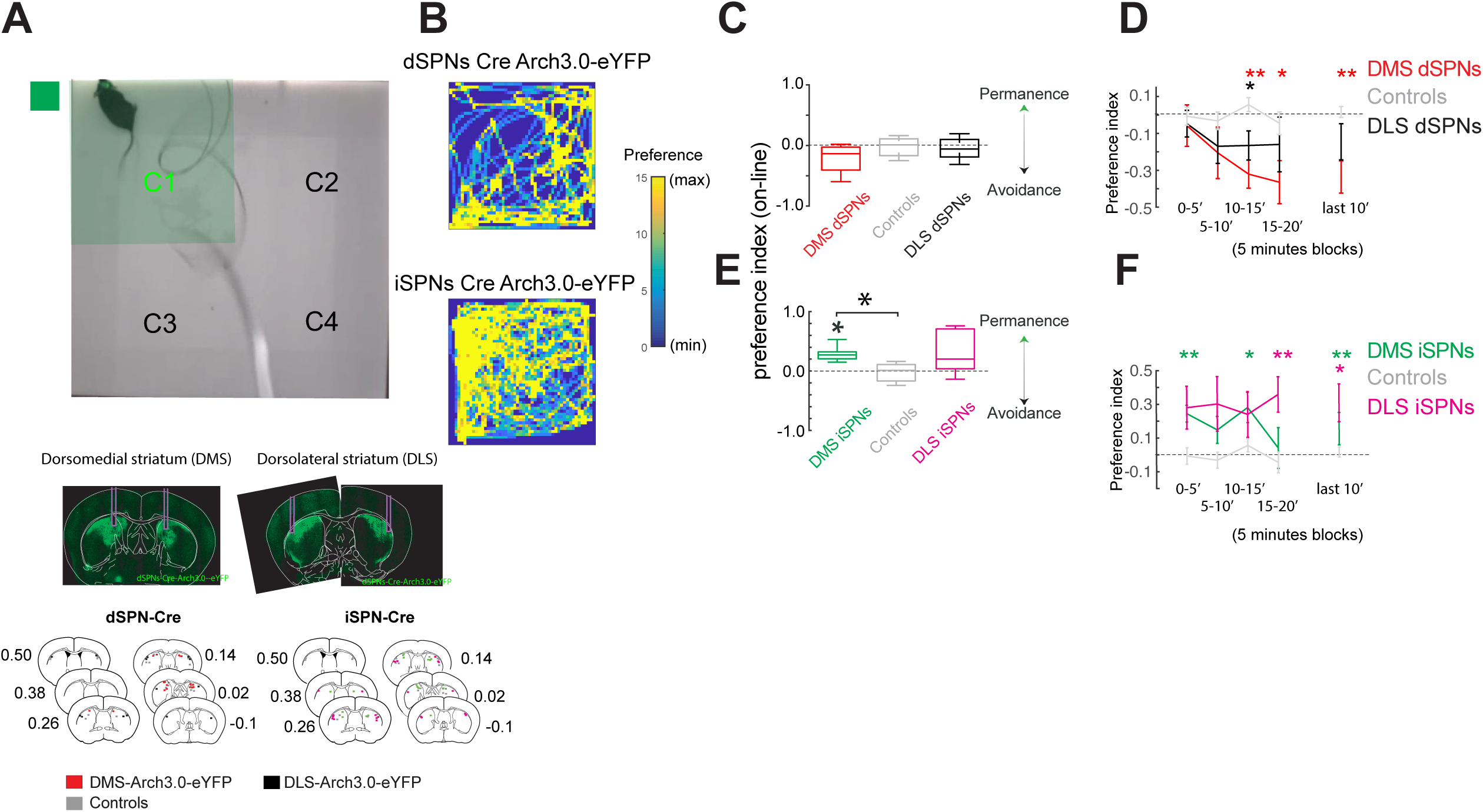
Optogenetic inhibition of dSPNs or iSPNs evokes avoidance or preference in the realtime place preference test. **A**. An open field was virtually divided into four quadrants, the quadrant where the animal received self-inhibition is highlighted in green. Middle panels show representative photomicrographs of coronal brain slices of animals with bilateral DMS or DLS implants. Bottom panels, schematic representation of the position of the fibers tips. **B**. Example of the trajectories of two different animals, color coding the time spent per pixel during the test. Inhibition of dSPNs or the iSPNs is shown in the upper and lower panels respectively. **C**. Quantification of the time spent in the quadrant with inhibition compared to the average of the other three quadrants. **D**. Same data as in C but plotted in bins of 5 min and the last 10 minutes in the test. **E**. and **F**. Same as C and D for animals with manipulation of iSPNs in the DMS, DLS and Controls. In E, * above the horizontal lines is the Dunńs multiple comparison test p<0.001, post Kruskal Wallis. * above a group is p<0.05 Wilcoxon test. In D and F, * above are p<0.05 Mann Whitney U test vs. the control group.

### Self-modulation of dSPNs and iSPNs during action selection supports opposite and complementary contributions of these pathways in the DMS and DLS respectively

To evaluate the effects of activity modulation of the striatal pathways on action selection (beyond approaching or avoiding a place), we performed a second experiment. In this experiment, animals had the option of selecting one of two actions: pressing a lever on the right or the left side of a food port. Either of the actions delivered a pellet but only one was paired with light delivery at the time of pressing (animals were counterbalanced) (***Figure 3A***). Animals that received self-stimulation of dSPNs, either in the DMS (n=8) or DLS (n=9), preferred to select the lever paired with self-stimulation compared to controls [DLS-D1-ChR2 vs. controls (n=20), RM-ANOVA of the last 4 days of acquisition, group effect: F(1,25)=7.13, p=0.01; DMS-D1-ChR2 vs. controls, group effect: F(1,27) =5.33, p=0.02, ***Figure 3B***, left panel]. The opposite was observed when animals received self-stimulation of iSPNs, both in the DMS (n=7) and DLS (n=10) (RM-ANOVA of the last 4 days of acquisition, DLS-A2A-ChR2 vs. controls (n=20), group effect: F(1,28)=6.15, p=0.01; DMS-A2A-ChR2 vs. controls, group effect: F(1,25)=7.42, p=0.01, ***Figure 3C***, left panel).

**Figure 3.**
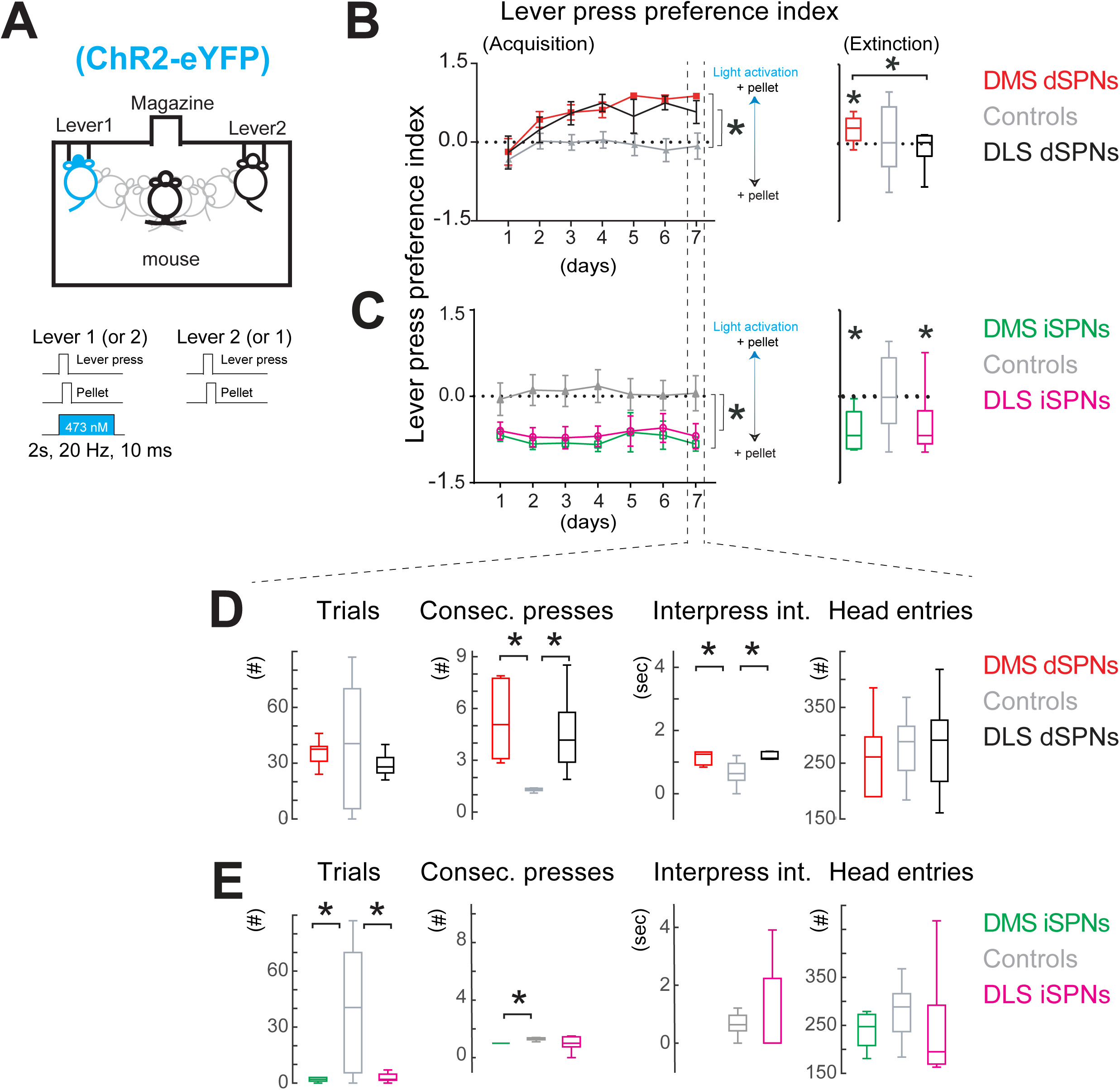
Self-activation of striatal dSPNs or iSPNs in an action selection test. **A**. Schematic of the task: operant chamber with two levers extended, at the moment of executing the action, pressing any of the levers delivered a pellet but only one of them was paired to self-activation. Bottom panels: details of the pellet and light activation. **B**. Preference index of lever presses across acquisition sessions, comparing the self-activation of dSPNs in the DMS (n=8) vs. DLS (n=9) and Controls (n=20). **C**. Same as B, comparing self-activation of iSPNs in the DMS (n=7) vs. DLS (n=10) and Controls (n=20). Right panel in B and C: preference index of the corresponding animals in an extinction test (here the actions yield no pellet and no light). D and **E**. Quantification of the behavioral execution of the actions on the lever that delivered self-activation on the last day of acquisition for animals that received self-activation of dSPNs or iSPNs, respectively. * in B and C left panels: p<0.05 RM-ANOVA. In B, C, right panels and D and E, * above the horizontal lines is the Dunńs multiple comparison test p<0.05, post Kruskal Wallis. * above a group is p<0.05 Wilcoxon test.

Analysis of presses on the lever paired with light delivery, during the last day of acquisition, revealed that the activation of dSPNs, either in the DMS or DLS, increased the number of actions (mean±SEM, consecutive presses: DMS-D1-ChR2=5.3±0.9; DLS-D1-ChR2=4.5±0.8; Controls=1.6±0.4, p<0.05, Kruskal Wallis test, ***Figure 3D***, column 2), but intriguingly slowed down the inter-press intervals (DMS-D1-ChR2=1.3±0.2; DLS-D1-ChR2=1.2±0.1; Controls=1.7±1.1, p<0.05, Kruskal Wallis test, ***Figure 3D***, column 3). Activation of iSPNs, either in the DMS or DLS, decreased the number of trials (mean±SEM, times that an animal pressed the lever and then visited the food port; DMS-A2A-ChR2=5±3; DLS-A2A-ChR2=4±1.6; Controls=39±9, p<0.05, Kruskal Wallis test, ***Figure 3E***, column 1), but interestingly, consecutive presses were only decreased when activating this pathway in the DMS (mean±SEM, DMS-A2A-ChR2=0.8±0.17; DLS-A2A-ChR2=0.9±0.19; Controls=1.69±0.49, p<0.05, Kruskal Wallis test, ***Figure 3E***, column 2).

To evaluate whether self-activation during acquisition has lasting effects on action-outcome memory we measured the action selected by animals during an extinction test (in the absence of either pellets or light modulation). Only animals that had previously received activation of dSPNs in the DMS, but not in the DLS, reported the same preference as during the acquisition [mean±SEM ratio of lever presses, DMS-D1-ChR2=0.24±0.08, p<0.028, Wilcoxon test, Figure 3B, right panel]. However, animals that had previously received self-stimulation of iSPNs, either in the DMS or the DLS, avoided the lever previously paired with light activation [DMS-A2A-ChR2=-0.59±0.12, A2A-ChR2-DLS=-0.50±0.14, p<0.029, Wilcoxon test, ***Figure 3C***, right panel]. This extinction test shows that in the DMS the striatal pathways have opposite contributions to the selection of actions but in the DLS they do not. This suggests a difference in the memory of mice depending on whether such memory was acquired with the activation of dSPNs in the DMS versus the DLS.

To get a clearer picture of the contribution of the striatal pathways to action selection we performed the inhibition experiment, this time pairing the pressing of one of the levers with inhibition of the dSPNs or iSPNs in the DMS vs. DLS (***Figure 4A***). Animals that received self-inhibition of dSPNs in the DMS avoided pressing the lever paired with self-inhibition [DMS-D1-Arch3.0 (n=8; 2 D1-Cre/Ai35) vs. controls (n=17), RM-ANOVA, group effect: F(1,23)=5.19, p=0.03, ***Figure 4B***, red line). Surprisingly, animals that received self-inhibition of dSPNs in the DLS selected the action paired to self-inhibition (DLS-A2A-Arch3.0 (n=6; 2 D1-Cre/Ai35) vs. controls (n=17), RM-ANOVA, group effect: F(1,21)=5.69, p=0.02, ***Figure 4B***, black line). On the other hand, animals that received self-inhibition of iSPNs, either in the DMS or the DLS, selected the action paired with self-inhibition [DMS-A2A-Arch3.0 (n=8; 3 A2A-Cre/Ai35) vs. controls (n=17), RM-ANOVA, group effect: F(1,23)= 4.28, p= 0.04, ***Figure 4C***, green line; DLS-A2A-Arch3.0 (n=8; 4 A2A-Cre/Ai35) vs. controls (n=17), group effect: F(1,23)=4.40, p=0.04, ***Figure 4C***, pink line]. This experiment again revealed that the striatal pathways in the DMS have opposite contributions, but that in the DLS, inhibition of dSPNs or iSPNs yielded a similar effect suggesting a complementary contribution of the pathways in this compartment.

**Figure 4.**
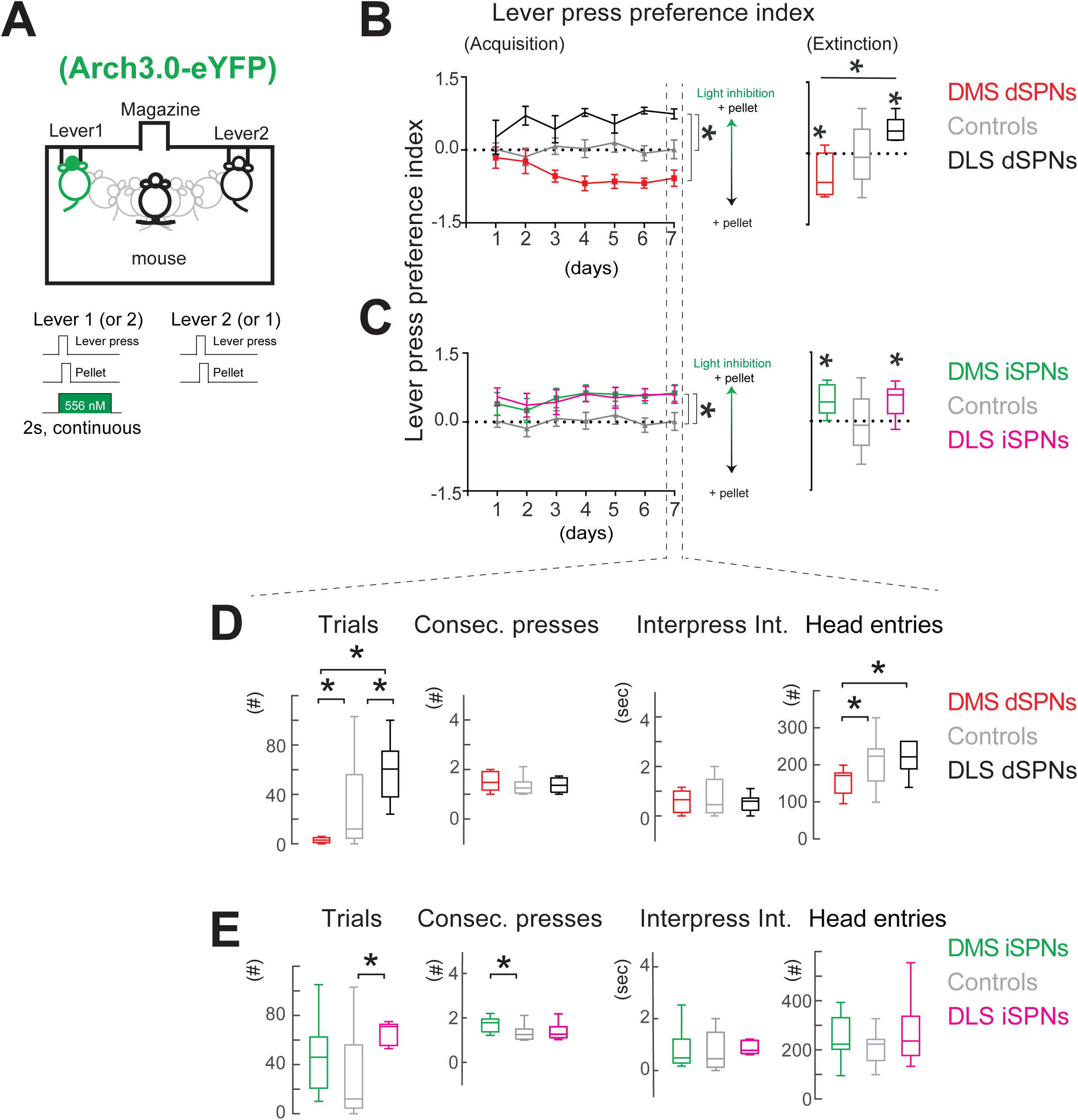
Self-inhibition of striatal dSPNs or iSPNs in an action selection test. **A**. Schematic of the task: operant chamber with two levers extended, at the moment of executing the action, pressing any of the levers delivered a pellet but only one of them was paired to self-inhibition. Bottom panels: details of the pellet and light inhibition. **B**. Preference index of lever presses across acquisition sessions, comparing the self-inhibition of dSPNs in the DMS (n=8) vs. DLS (n=6) and Controls (n=17). **C**. Same as B, comparing self-inhibition of iSPNs in the DMS (n=8) vs. DLS (n=8) and Controls (n=17). Right panel in B and C: preference index of the corresponding animals in an extinction test (here the actions yield no pellet and no light). **D** and **E**. Quantification of the behavioral execution of the actions on the lever that delivered self-inhibition on the last day of acquisition for animals that received self-inhibition of dSPNs or iSPNs, respectively. * in B and C left panels: p<0.05 RM-ANOVA. In B, C, right panels and D and E, * above the horizontal lines is the Dunńs multiple comparison test p<0.05, post Kruskal Wallis. * above a group is p<0.05 Wilcoxon test.

Analysis of presses on the lever paired with the delivery of light inhibition, during the last day of acquisition, revealed that only the inhibition of dSPNs in the DMS decreased the number of trials (mean±SEM, DMS-D1-Arch3.0=3±1; Controls=28±7, p<0.003, Kruskal Wallis test, ***Figure 4D***, column 1). Remarkably, these effects were the opposite to those seen when inhibiting dSPNs in the DLS (DMS-D1-Arch3.0 vs. DLS-D1-Arch3.0=59±11, p<0.05, Mann Whitney U test, ***Figure 4D***, column 1). A similar result was observed when comparing head entries (***Figure 4D***, column4), implying a difference in the contribution of dSPNs to the selection of actions in the DMS versus the DLS. On the other hand, inhibition of iSPNs in the DMS and DLS increased the execution of actions either by increasing the number of trials (DLS-A2A-Arch3.0=59±8 vs. controls= 28±7, p<0.05, Kruskal Wallis test, ***Figure 4E***, column 1) or the consecutive presses (DMS-A2A-Arch3.0=1.6±0.1 versus Controls=1.2±0.1, p<0.05, Kruskal Wallis test, ***Figure 4E***, column 2) without affecting the interpress intervals or head entries (***Figure 4E***, columns 3-4).

Notably, when evaluating the effect of the inhibition in the extinction test, animals selected the same action as during the acquisition [mean±SEM ratio of lever presses DMS-D1-Arch3.0=- 0.47±0.15 (n=8; 2 D1-Cre/Ai35), DLS-D1-Arch3.0=0.51±0.10 (n=6; 2 D1-Cre/Ai35), DMS-A2A-Arch3.0= 0.45±0.12 (n=8; 3 A2A-Cre/Ai35), DLS-A2A-Arch3.0= 0.43±0.14 (n=7; 4 A2A-Cre/Ai35), p<0.05, Wilcoxon test, ***Figures 4B*** and ***4C***, right panels).

Taken together, the results from modulating the activity of dSPNs or iSPNs during action selection revealed that an opposite contribution of striatal pathways holds for the DMS, but that for the DLS, a complementary contribution better supports this behavior.

### Context-dependent contribution of the striatal pathways to reward-seeking versus spontaneous displacements

Finally, to evaluate the contribution of striatal cells to the displacement of animals we evaluated the effects of activating or inhibiting dSPNs or iSPNs in the DMS versus the DLS in two contexts: when displacing to collect a pellet, in the previously described lever press task (reward seeking displacements guided to reach a palatable goal) and in an open field (spontaneous displacements without a palatable goal) (***Figures 5-6***).

**Figure 5.**
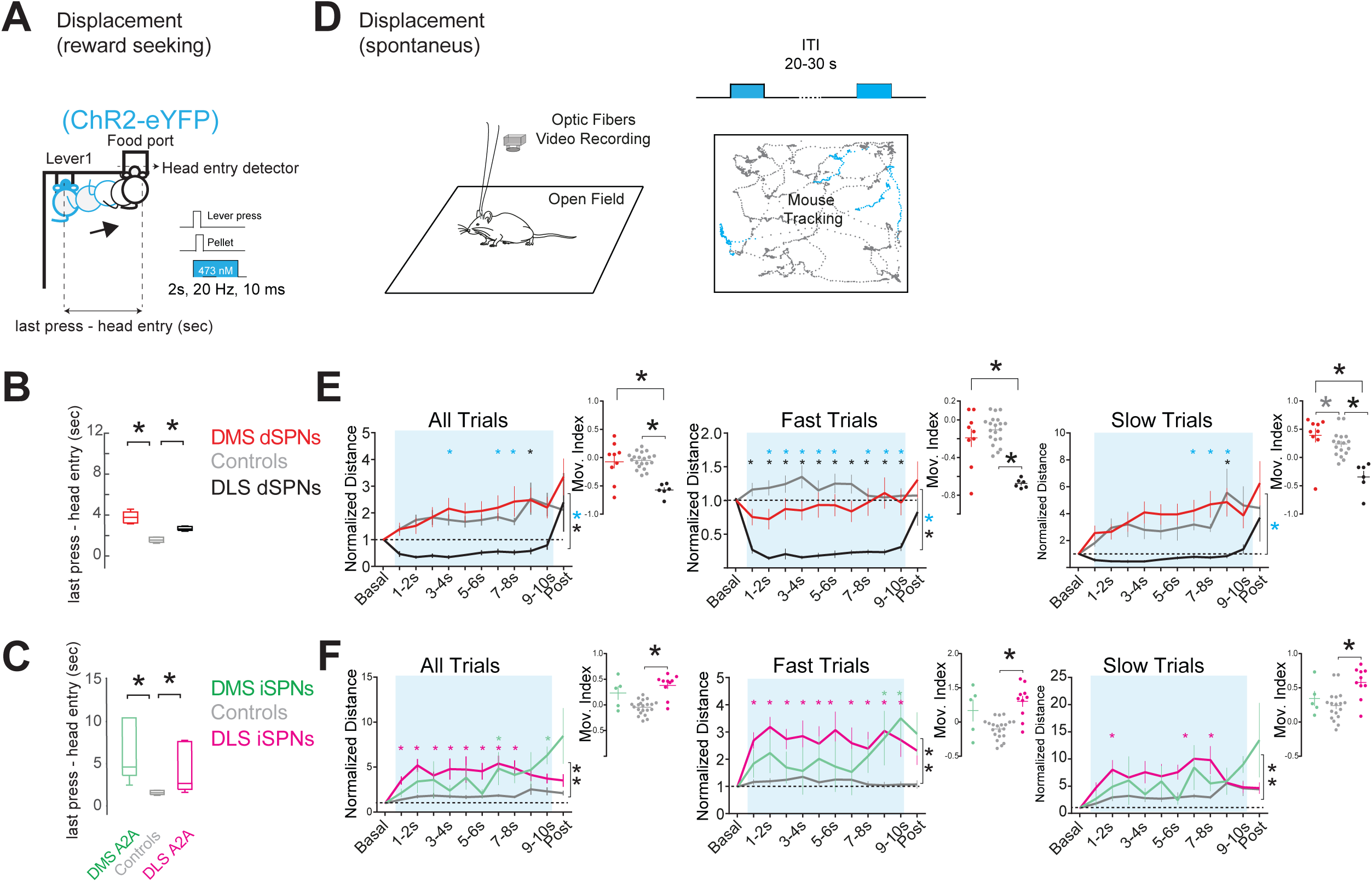
Activation of striatal dSPNs or iSPNs during reward seeking and spontaneous displacements in the open field. **A**. Schematic of the displacement of animals from the lever that delivered self-activation in the same setup as Figures 3 and 4. In this case, since this is the last day of acquisition of the action (lever press) that delivered an outcome (pellet) animals developed stereotyped displacements from the lever to the food port. Light modulation was switched on at the initiation of the action. **B** and **C**. Time from executing the action to visiting the food port for animals that received activation of dSPN or iSPNs, respectively. **D**. Schematic of the activation in the open field, in which light was delivered every 20-30 sec, with the same protocol as during the operant task (20 Hz, 10 ms pulses), right panel: schematic of the tracking of an animal. **E**. Normalized distance of animals with activation of dSPNs in the DMS or DLS and Controls. Trials (All trials) were categorized as fast or slow trials depending on the mean displacement of the animal (one second) before the activation (see methods). **F**. Same as E, for animals that received iSPNs activation in the DMS DLS and Controls. In B, C and the right panels of E and F, * above the horizontal brackets is the Dunńs multiple comparison test p<0.05, post Kruskal Wallis, except in E right panel slow trials, where the grey * is p<0.05 Mann Whitney U test. * beside the vertical brackets in the left panels of E and F is p<0.05, RM-ANOVA, and above are p<0.05, Sidak’s multiple comparison, blue asterisks depict the DMS vs. DLS comparison.

**Figure 6.**
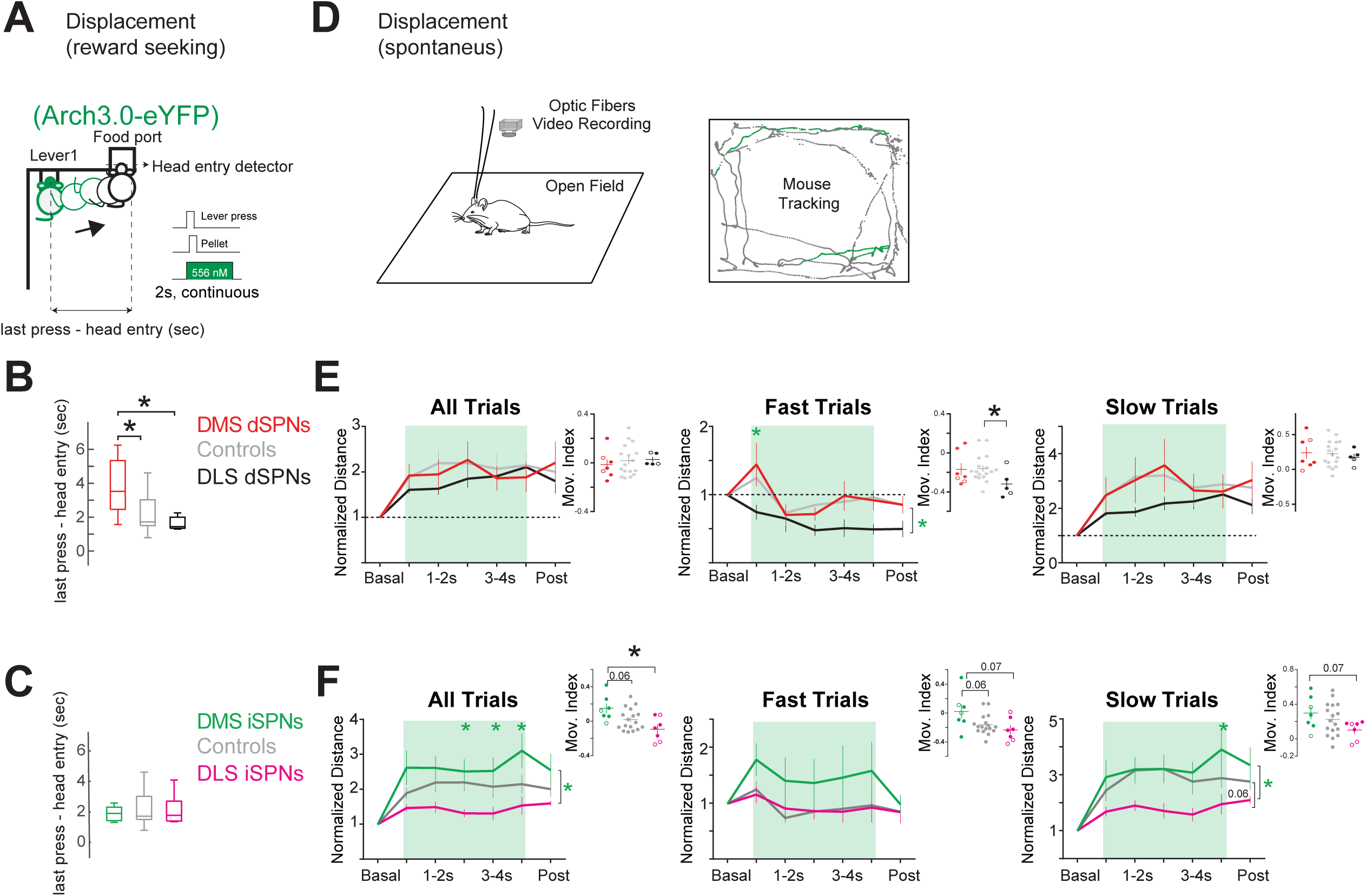
Inhibition of striatal dSPNs or iSPNs on reward seeking and spontaneous displacements in the open field. **A**. Schematic of the displacement of animals from the lever that delivered self-inhibition in the same setup as figures 3 and 4. In this case, since this is the last day of acquisition of the action (lever press) that delivered an outcome (pellet) animals developed stereotyped displacements from the lever to the food port. Light modulation was switched on at the initiation of the action. **B** and **C**. Time from executing the action to visiting the food port for animals that received inhibition of dSPN or iSPNs, respectively. **D**. Schematic of the inhibition in the open-field, in which light was delivered every 20-30 sec, with the same protocol as during the operant task (2-second continuous pulse), right panel: schematic of the tracking of an animal. **E**. Normalized distance of animals with inhibition of dSPNs in the DMS (n=7) or DLS (n=5) and Controls (n=17). Trials (All trials) were categorized as fast or slow trials depending on the mean displacement of the animal (one second) before the activation (see methods). **F**. Same as E, for animals that received inhibition of the iSPNs [DMS (n=7); DLS (n=7); Controls (n=17)]. In B and F, * above the horizontal brackets is the Dunńs multiple comparison test p<0.05, post Kruskal Wallis. * in E right panel fast trials is p<0.05 Mann Whitney U test. * beside the vertical brackets in E and F left panels is p<0.05, RM-ANOVA, and above are p<0.05, Sidak’s multiple comparison, green asterisks depict the comparison DMS vs. DLS.

Activation of the striatal pathways during reward seeking displacements i.e., modulating their activity when coming from a lever press to the food port (***Figure 5A***) revealed that activation of dSPNs or iSPNs, in either the DMS or DLS, slowed the displacement of animals and increased the time taken to reach the food port [(median) mean±SEM: DMS-D1-ChR2=(3.8) 3.8±0.2; DLS-D1-ChR2=(2.6) 3±0.3; Controls=(1.5) 3.4±1.3, p<0.05, Kruskal Wallis test, ***Figure 5B***; DMS-A2A-ChR2=(4.6) 8.6±4.6; DLS-A2A-ChR2=(2.6) 6.1±2.4; Controls=(1.5) 3.4±1.3, p<0.05, Kruskal Wallis test, ***Figure 5C***]. However, activation during spontaneous displacements in the open field (***Figure 5D***) had a different effect. Trials when animals were displacing were separated into fast and slow trials (see methods and Ramirez-Armenta et al., 2022). Activation of dSPNs in the DMS increased the displacement of animals in slow trials [***Figure 5E*** slow trials: mean±SEM, movement index DMS D1-ChR2 =0.39±0.12 (n=9) vs. Controls=0.24±0.05 (n=12), p<0.05, Mann Whitney U Test], whereas activation of dSPNs in the DLS decreased the displacements of all, fast and slow trials [***Figure 5E***, all and fast trials, normalized distance DLS-D1-ChR2 (n=6) vs Controls (n=12), RM-ANOVA, p<0.05; movement index all, fast and slow trials, p<0.001, Kruskal Wallis Test]. On the other hand, activation of iSPNs in either the DMS or the DLS increased displacement in all, fast or slow trials [***Figure 5F***: normalized distance DMS-A2A-ChR2 (n=5) or DLS-A2A-ChR2 all (n=10) vs. Controls (grey, n=21), RM-ANOVA, p<0.05]; particularly when activating these cells in the DLS versus Controls (movement index, p<0.001, Kruskal Wallis Test).

The inhibition experiment showed that on reward seeking displacements only the inhibition of the dSPNs in the DMS slowed the displacement of animals [more time taken to reach the food port: (median) mean±SEM, DMS-D1-ChR2=(3.5) 3.8±0.6 (n=7; 2 D1-Cre/Ai35) vs. Controls=(1.7) 2.3±0.4 (n=17), p<0.05, Kruskal Wallis test, ***Figure 6A-B***], with iSPNs inhibition interestingly having no effect in this context (***Figure 6C***). Intriguingly during spontaneous displacements in the open field (***Figure 6D***), the inhibition of dSPNs in the DLS decreased the fast trials (***Figure 6E***, movement Index panel: DLS-D1-Arch3.0, fast trials=-0.32±0.06 (black, n=5; 2 D1-Cre/Ai35), vs. Controls fast trials=-0.16±0.03 (grey, n=17), p<0.05, Mann Whitney U Test). In this second context the inhibition of iSPNs in the DMS vs. the DLS showed opposite effects [***Figure 6F***, normalized distance DMS-A2A-Arch3.0 in all trials or slow trials (n=7; 2 A2A-Cre/Ai35) vs. A2A-DLS-Arch3.0 (n=7; 3 A2A-Cre/Ai35), RM-ANOVA, p<0.05] with a tendency for the inhibition of iSPNs in the DMS to increase locomotion (***Figure 6F***, movement Index panel: DMS-A2A-Arch3.0 all trials=0.15±0.06 vs. Controls all trials =0.019±0.03 (n=17), p=0.06, Mann Whitney U Test). Interestingly, inhibition of the iSPNs in the DLS showed a tendency to decrease locomotion exclusively in slow trials (***Figure 6F***, slow trials normalized distance panel, Two-way ANOVA, p=0.06).

Together the results from manipulating the activity of striatal SPNs during displacement in two different contexts revealed that on reward seeking displacements the activity of the dSPNs in the DMS is required for these displacements (but that this is not the case for dSPNs in the DLS, or iSPNs in the DMS or DLS). However, when executing spontaneous displacements in an open field, boosting the activity of dSPNs in the DMS can increase displacements, although in this second context, the activity of the dSPNs and iSPNs in the DLS showed to be driver of spontaneous displacements.

## Discussion

The results presented here show that the activity of the SPNs of the two striatal pathways have specific contributions to the behavior of animals depending on the striatal compartment and on the demands of the context. The main findings of this study are: (1) in real-time place preference self-modulation of dSPNs or iSPNs, either in the DMS or DLS, has opposing contributions, with dSPNs supporting preference and iSPNs avoidance (***Figure 7A***, column 2). (2) in action selection, pairing the action to self-activation of dSPNs or iSPNs during acquisition of the action-outcome relationship, either in the DMS or DLS, leads to selecting or avoiding the action associated with self-activation. However, pairing the action with self-inhibition, resulted in opposing effects of the pathways only in the DMS; in the DLS the pathways had a complementary contribution (***Figure 7A***, column 3). (3) During locomotion to reach a goal, activation of either of the pathways in the DMS or DLS slowed displacements, as did inhibiting the direct pathway in the DMS (***Figure 7A***, column 5). Conversely, during spontaneous displacements without a palatable goal, activation of dSPNs or inhibition of iSPNs in the DMS increased locomotion, though inhibition of dSPNs or activation of iSPNs in the DMS has no effect or increased spontaneous displacements respectively; remarkably, inhibition of either of the pathways in the DLS impaired spontaneous displacements (***Figure 7A***, column 4). Overall, our findings suggest that the the activity of the of striatal pathways in the DMS vs. DLS support the model of opposite contributions in real-time place preference but in action selection or locomotion either a complementary or an undescribed model is better supported (Figure 7B).

**Figure 7.**
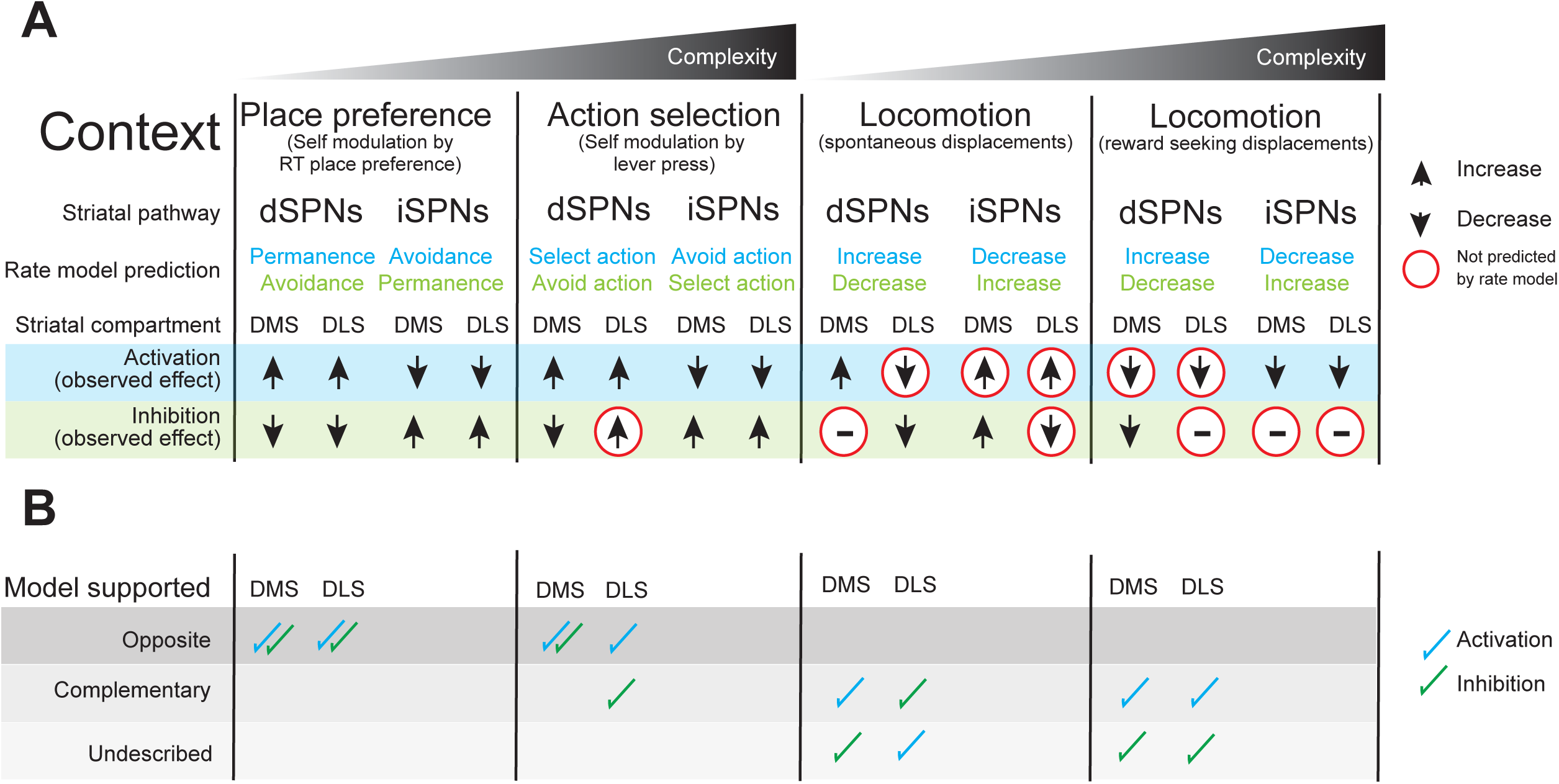
Context-dependent contributions of the striatal pathways in the associative and sensorimotor striatum. **A**. Summary of the findings of activation and inhibition experiments according to striatal compartment. **B**. Model supported by the presented findings in each striatal compartment. Opposite: modulation of the striatal pathways has opposite effects on behavior. Complementary: modulation of the striatal pathways has similar effects. Undescribed: neither of the models was observed.

### Do ChR2- and Arch3.0-mediated modulations of SPNs yield activation and inhibition of neurons respectively?

We and other groups have previously documented the feasibility of using ChR2 or Arch3.0 expressed in SPNs to activate or inhibit respectively these neurons in the DLS (Tecuapetla et al., 2014, 2016) or the DMS (Kravitz et al., 2010; Tai et al., 2012; Ramírez-Armenta et al., 2022; Bolkan et al., 2022). We have also reported that activation of ChR2 above 14 Hz may induce inactivation blocks in a small percentage of cells, although the population effect is activation (Tecuapetla et al., 2016). Therefore in the present study, we aimed to compare the result from every activation experiment with that of the corresponding inhibition experiment. Furthermore, since heating tissue in the absence of opsins may have effects (Owen et al., 2019), every experiment was compared to control animals expressing only eYFP in the absence of opsins and undergoing the same procedure.

### Why does self-modulation of dSPNs in the DLS have complimentary effects on complex action selection?

Self-activation of the striatal pathways, in the real-time place preference test and when selecting a lever in the action selection test demonstrated opposing contributions to the tasks. However, self-inhibition of the pathways supported opposing contributions only in the DMS, with the DLS showing complementary contributions. A possible explanation is that the demands of these tests are different. While the online place preference test requires displacing the body of the animal, the motivation to displace comes from the effects of the striatal pathway’s modulation. However, in the action selection test the motivation comes from the intrinsic mechanisms to get the goal (pellet) plus the modulation of the pathways. The fact that self-modulation of the DMS yielded similar effects in the place preference and action selection tests strongly argues that the contribution of the striatal pathways in this compartment is similar independent of the action selection test, whether it implies selecting where to move the body or where to execute a more explicit action. On the other hand, the results of the self-modulation of striatal pathways in the DLS strongly suggests that in this compartment the opposition model of the pathways does not hold. Instead, since in the DLS inhibition of either pathway yielded an increase in the selection of the action a complementary contribution is supported. There is also a further question: Why did the activation experiments support opposing contributions of the pathways both in the DMS and DLS whilst the inhibition experiments did this only in the DMS but not in the DLS? One explanation is that as previously mentioned, for the place preference test the main driver was the self-modulation of the pathways, but in the action selection test the intrinsic activity leading the searching for the goal plus the optogenetic modulation of the pathways occurred. We speculate that the optogenetic self-activation can overcome the intrinsic activity both in the place preference and in the action selection test (suggesting that the artificial activation of the striatal pathways may impose effects). However, when it comes to optogenetic inhibition, particularly for the striatum which is a structure of low baseline activity (Jin et al., 2014), it relies more on removing the intrinsic activity in the different contexts. Therefore, in the experiment pairing self-inhibition with the selection/execution of the action, inhibiting dSPNs in the DLS was not enough to block the selection of the action unlike in the DMS. In fact, the inhibition of it in the DLS favored the execution of the action, perhaps silencing a mechanism that normally would be opposed to the selection of actions.

### Why does the modulation of pathway activity yield differential contributions to reward seeking and spontaneous displacements?

Previously it has been shown that dSPNs activation in the DMS speeds up the displacement of animals (in an open field) or the latency to execute actions (in a nose poke two-choice task), while the activation of the iMSNs shows opposite effects (Kravitz et al., 2010; Tai et al., 2012). Also, it has been shown that activation of either of the striatal pathways in the DLS pauses or increases the latency to execute actions [in a single lever fixed ratio-8 task; (Tecuapetla et al., 2016)]. However, a direct comparison between the activation of the striatal pathways in the DMS vs. the DLS had not been performed. In this study, we aimed to compare the effects of the optogenetic modulation of the striatal pathways in two different contexts, in displacements after selecting an action that leads to an outcome (pellet), and on spontaneous displacements without an explicit goal (in the open field). During the displacement to reach a goal, the activation of either of the striatal pathways in the DMS or DLS slowed locomotion and increased the time from executing the action to visiting the food port (***Figure 5A-C***). According to the rate model of the striatal pathways (Albin et al., 1989; DeLong, 1990; Penney and Young, 1983), it was expected that activation of dSPNs and iSPNs would favor and slow displacements respectively. The latter was consistent with our experimental findings, but the former was not. One explanation for the discrepancy between the expected and the experimental findings of the activation of the direct pathway SPNs could be that the 20-Hz activation used in this study interfered with or blocked the natural spiking of some of these cells. To investigate this possibility we performed additional experiments, this time inhibiting the striatal pathways in the same context. It was expected that inhibition of dSPNs and iSPNs would slow and speed up displacements respectively (Albin et al., 1989; DeLong, 1990; Penney and Young, 1983; Kravitz et al., 2010). Experimentally, we observed that only the inhibition of dSPNs in the DMS slowed displacements. Together the results from the activation and inhibition experiments in this context allow us to conclude that in the DMS dSPNs activity supports displacing to reach a goal. However, although activation of iSPNs slowed down displacements to reach a goal, the fact that their inhibition did not speed them up, means that we cannot conclude that iSPNs activity supports the reaching of a goal.

During spontaneous displacements without an explicit goal, the activation of dSPNs in the DMS increased locomotion (***Figure 5E***, slow trials) as previously reported (Kravitz et al., 2010), albeit in a specific set of trials. Conversely, activation of dSPNs in the DLS decreased all types of displacements in the open field (***Figure 5E***), consistent with a report showing that high dSPNs activation in the DLS pauses the displacing of animals (Tecuapetla et al., 2016). In the case of the indirect pathway, its activation both in the DMS and DLS increased spontaneous displacements, an unexpected finding in the DMS (Kravitz et al., 2010) but not for the DLS (Tecuapetla et al., 2016). Since findings from the activation of the striatal pathways may have been caused by interrupting the natural spiking of the SPNs, we also performed the complimentary inhibition experiments. Intriguingly, inhibition of dSPNs in the DLS, but not the DMS, decreased spontaneous displacements while inhibition of the iSPNs in the DMS resulted in an increase compared to inhibition in the DLS in this context. From this set of findings, we conclude that during spontaneous displacements without a goal, there is an activation range for dSPNs in the DMS to increase locomotion, but that either interfering with or inhibiting the activity of this pathway in the DLS decreases locomotion in this context. Still, an open question is: Why during reward seeking displacements is it that dSPNs inhibition in the DMS reveals its participation whereas on spontaneous displacements in the open field it is activation that reveals its contribution in favoring displacements? We speculate that when reaching a goal, the dSPNs of the DMS may have a specific activity to reach the goal (interfering with this pattern would slow displacements as in ***Figure 5B***), while in spontaneous slow trial displacements, these cells may be silent, and boost locomotion when they are activated. On the other hand, dSPNs in the DLS may not contribute heavily to reaching a goal because their inhibition did not impair the displacement to reach the goal. Conversely, during spontaneous displacements this pathway in the DLS may have a level/pattern of activity that if interfered with (***Figure 5E***) or inhibited (***Figure 6E***, fast trials) it decreases locomotion. Finally, during spontaneous displacements, increasing indirect pathway activity in the DMS or DLS increased locomotion (although we do not rule out that our activation of this pathway in this context may have interfered with natural spiking; although note that this same modulation was used in the place preference and acquisition of lever press where canonical findings were observed), and the fact that only its inhibition in the DMS increased locomotion allowed us to conclude that this pathway in the DMS supports the control of spontaneous displacements, not necessarily by signaling motor suppression but aversion (Isett et al., 2023).

## Conclusions

In the present study self-modulation of the striatal pathways, during real-time place preference or action selections, in the DMS supported opposing contributions whereas in the DLS opposing or complementary contributions (***Figure 7A-B***, columns 2-3). When testing the contribution of the pathways to locomotion, they showed differential contributions depending on the striatal compartment and on the context, which does not always support either an opposite or a complementary model (***Figure 7A-B***, columns 4-5). Our data suggest that in future studies of striatal function, both compartments and contexts must be considered.

## Materials and Methods

### Animals

We crossed BAC transgenic mice expressing Cre recombinase under control of the D1 (D1-Cre EY217) or A2A (A2A-Cre KG139) promoter with wild-type C57BL/6J or Ai35 (Jax: 012735) mice for at least six generations. Heterozygotous male and female mice between 8 and 12 weeks of age were used for experiments and by food restrictions maintained at 80-85% of their original weight. Animals were housed in a 12/12 h light/dark cycle. On postnatal day 21, tail tissue samples were collected for DNA extraction and genotyping. DNA extractions were performed with a kit (F140 WH, Thermo Scientific), after which Cre-recombinase was amplified by PCR. All animal procedures were performed following the animal care committee regulations from the Instituto de Fisiologia Celular of the Universidad Nacional Autónoma de México (FTA-121-17).

### Stereotaxic surgery: virus injections and fiber implantation

A viral vector was used to express channelrodhopsin (AAV1.EF1a.DIO.hChR2(H134R)-eYFP.WPRE.hGH; UNC Vector core), or the reporter protein eYFP in control animals. For inhibition experiments Ar-chaerhodopsin3.0 (UNC Vector Core), D1-Cre/Ai35 or A2A-Cre/Ai35 mice were used. Anesthesia was induced with 4% isofluorane and animals were maintained under anesthesia with 1% isofluorane and oxygen (1 L/min) using an anesthesia system (E-Z Anesthesia) coupled to the stereotaxic apparatus (David Kopf Instruments). Thermal support was provided with a heat pad (Physitemp Instruments). After aseptic procedures (ethanol 70% and iodine solution), hair was removed from the cephalic zone, and the skull was exposed and cleaned. The skull was then aligned and a bilateral trephination was performed using a dental drill (marathon SDE-H355P1) with the following coordinates: for DMS +0.5 mm anteroposterior (AP), ±1.6 mm mediolateral (ML), −2.3 mm dorsoventral (DV); for DLS +0.5mm AP, ±2.5 mm ML, −2.3 mm DV. Bregma was used as AP and ML reference and the surface of the brain as the DV reference. Virus was injected bilaterally in the DMS or DLS at 4.6 nL/5 seconds (9 minutes) using a borosilicate pipette (Drummond scientific, >25 µm) coupled to a nanoinjector (Nanoject II Drummond Scientific). Pipettes were kept in the brain for 15 minutes after injections to allow virus diffusion. After bilateral injections, fiber optics of 200 µm or 300 µm in diameter were implanted for excitation or inhibition experiments respectively. Fiber implants were custom-made following the protocol of (Sparta et al., 2012). Finally, we covered the skull with dental acrylic (Ortho jet) and then supervised the recovery of animals for 4-7 days.

### Real-Time Place Preference Test

Animals were placed into an open box (40 cm wide x 40 cm long x 31 cm high) for 15-20 minutes. A quadrant was selected at random, and animals received stimulation or inhibition when they moved inside this quadrant (Bonsai Software). Videos were recorded from above at 30 fps and the body of animals was tracked using Bonsai Software (Lopes et al., 2015). This allowed the time spent in the selected quadrant to be quantified.

### Training mice to lever press

Pre-training (session 1): animals received sugar pellets randomly for 30 minutes (14 mg Bio-Serv) in the reward port located in the center of the chamber (MED-Associates, Inc). Continuous reinforcement 15 (session two) and 30 (session three): a lever was extended on the right or left side of the reward port (counterbalanced between mice) and animals were rewarded with a pellet when they pressed the lever. Sessions ended after animals pressed the lever 15 times (CRF15) or 30 times (CRF30), or after 30 minutes had elapsed, whichever happened first. Next, animals were subjected to a CRF30 session with a second lever extended on the opposite side of the chamber. This second stage of training was the acquisition of the action-outcome of the two levers and consisted of 4-7 sessions. Sessions began with the illumination of the chamber and extension of both levers and finished with the retraction of both levers and the light turning off. Both actions were rewarded (i.e., pressing the left or the right lever) but only one side was paired with light manipulation. The lever with stimulation/inhibition was on the opposite side of the chamber to the lever extended in the last CRF30 training session. Extinction test: after the last day of the acquisition training, all animals received a 10-minute extinction test. Both levers were extended but animals did not receive rewards or self-stimulation after pressing. Event timestamps at a resolution of 10 milliseconds (lever press, head entries, simulations) and video were recorded for all sessions.

### Measuring locomotion guided by reaching a goal

The timestamps recorded above were used to calculate the number of trials (times that an animal performed presses and then a head entry into the reward port), consecutive presses (series of presses without a head entry), inter-press interval, number of head entries, and locomotion guided by reaching the goal (time between a head entry and the press that preceded it).

### Measuring locomotion without a goal in the open field

Animals were placed into an open box and video was recorded from above at a resolution of 30 fps. The body of animals was tracked using Bonsai Software to obtain centroid coordinates. Horizontal displacement of animals was calculated with custom Matlab scripts, as the sum of the Euclidean distances of centroid coordinates between frames. We used one-second time windows to define displacement. Each value was normalized to 1 sec before the start of each light pulse and this was used as the locomotion baseline. As described in a previous study (Ramírez-Armenta et al., 2022), we classified trials as fast or slow, using a fixed locomotion threshold related to the mean locomotion before the light pulse. We define the movement index as the ratio between the mean displacement during the light manipulation minus the displacement during the basal period and divided by the sum of these two variables.

### Anatomical Verification

After all experiments were finished, animals were deeply anesthetized with ketamine/xylazine (85%-15%) i.p. and perfused. An incision was made in the left ventricle of the heart and 15 mL of 0.1M PBS followed by 4% PFA (paraformaldehyde) was administered using a perfusion system. Brains were carefully extracted and kept in 4% PFA for 24 hours before being transferred to 0.1 M PBS. They were then embedded in 4% agar and 100 µm coronal slices were cut using a Vibratome (Series 1000, Ted Pella Inc). Slices were mounted on slides and stained with Hoescht (20 µg/mL). Anatomical verification was performed using an epifluorescence (Leica DM1000) or confocal microscope (Zeiss, LSM710).

### Data Analysis

MED-PC-IV software was used to count the timestamps of events in sessions in the MED PC boxes. For the place preference and open field tests, the centroid coordinates of animals were obtained through the use of bonsai and Matlab custom scripts. Each data analysis is specified in the text and figures. Statistical analyses were performed in GraphPad Prism and Matlab.

## Acknowledgments

We thank Professor Rui Costa for the D1-Cre and A2A-Cre mice and PhD Catherine French for proofreading the English of the manuscript.

## Author contributions

Cuevas and Tecuapetla wrote the manuscript. Tecuapetla, Cuevas, and Llanos designed the experiments. Cuevas and Llanos performed all the activation and inhibition experiments respectively. Ramirez-Armenta developed the analysis pipeline for open-field experiments and generated the figures for the open-field. Alatriste-Leon contributed to the inhibition experiments. Ramirez-Jarquin performed all genotyping and maintenance of the transgenic lines used. All authors contributed to the review of the manuscript.

